# Cellular proliferation biases clonal lineage tracing and trajectory inference

**DOI:** 10.1101/2023.07.20.549801

**Authors:** Becca Bonham-Carter, Geoffrey Schiebinger

## Abstract

We identify a fundamental statistical phenomenon in single-cell time courses with clone-based lineage tracing. Through simple probabilistic arguments, we show how the relative growth rates of cells influence the probability that they will be sampled in clones observed across multiple time points. We support these arguments with a simple simulation study and a time-course of T-cell development, and we demonstrate that this bias can impact fate probability predictions from trajectory inference methods. Finally, we explore how to develop trajectory inference methods which are robust to this bias. In particular, we show how to extend *LineageOT* [1] to use data from clones observed across multiple time points.

## 1 Introduction

The development of high-throughput single-cell RNA-sequencing technologies has motivated a flurry of work on computational methods for inferring developmental trajectories from static snapshots [2]–[17] and, inter-connected and in parallel, methods for inferring cell lineage from DNA barcodes [18]–[25]. Developmental trajectories and cell lineage contain complementary information; having both data sources is valuable for trajectory inference [1], [18], [23], [26]–[31].

In this work, we identify a fundamental statistical bias that emerges from sampling cell lineage barcodes across a time course. We consider the setting in which cell state and lineage barcodes may be measured simultaneously (i.e., for the same cells) [20], [32], and copies of the same lineage barcodes may be observed over multiple time points (e.g., by repeated sampling from the same population). While these data have been collected and analyzed for some time [23], [25], [33], this bias has not yet been reported.

We focus on the case of static lineage barcodes, which are incorporated into the genome at an early stage of the developmental process, and then inherited by daughter cells but not modified over time. Note that lineage barcodes may not be detected in all cells that are sampled, resulting in barcoded datasets containing a subset of cells where cell state is known but the lineage barcode is not. Static lineage barcodes allow the tracing of *clones*: groups of cells that share a common barcode and therefore all share a common ancestor at the time of barcoding. If a clone is observed at only a single time point we will refer to it as a *single-time clone*, otherwise we will refer to it as a *multi-time clone*. The distinction between multi-time and single-time clones is important for understanding the source of the bias we discuss in this paper, and codifies clones by the level of information that they provide on developmental trajectories. Multi-time clones are particularly valuable for trajectory inference because they contain direct observations of developmental transitions. Indeed, recent methods including *CoSpar* [31] are designed to leverage multi-time clonal barcodes to infer cell fates.

We present a mathematical analysis that proves that the relative abundance of subpopulations is changed, or biased, in multi-time clonal datasets. The source of the bias is heterogeneous growth rates; cells with more descendants are more likely to be represented in multi-time clones. Therefore, more proliferative subpopulations are over-represented in multi-time clonal datasets. We prove the existence of this effect by simple analysis of probabilities, validating our arguments with proportions obtained from simulations as well as real data. We further show that the performance of trajectory inference methods such as *CoSpar*, which rely on this biased information, may be negatively impacted by the presence of this sampling bias.

Knowledge of this statistical phenomenon can help guide the design of robust trajectory inference methods. We introduce an extension of *LineageOT* [1], called *LineageOT-MT*, which incorporates information from multi-time clonal barcodes in addition to that from single-time clonal barcodes. We test the method on a barcoded time course of T-cell induction (see separate paper [34]), and also simulated datasets for which the ground-truth is accessible. We find that this extension offers a robust improvement over the original *LineageOT* and that it may be negatively impacted by the presence of the sampling bias. Comparing the performance of *CoSpar* and *LineageOT-MT* in the presence of the sampling bias, we provide recommendations for which method may be more robust in which settings.

The remainder of the paper is organised as follows. In Section 2 we present a mathematical derivation of the sampling bias. In Section 3 we demonstrate that the performance of trajectory inference methods may be negatively impacted by analyzing predictions from *Cospar* in the presence of this bias. In Section 4 we introduce *LineageOT-MT*, demonstrate improved performance over the original *LineageOT*, and repeat the analysis of *Cospar* in the presence of the sampling bias for *LineageOT-MT*. We conclude in Section 5 with a Discussion, where we explore how state-of-the-art methods for lineage-informed trajectory inference may be developed to overcome the impact of this sampling bias. Given the potential for the application of trajectory inference results to biomedical technologies and treatments, understanding and improving the accuracy of these methods is of critical importance.

## 2 Cell growth introduces bias in multi-time clonal barcodes

In this section, we present a derivation of the statistical sampling bias in multi-time clone data.

Consider a population of cells comprised of two subpopulations denoted type A and type B, where type B cells have a higher rate of proliferation than those of type A. Each cell is given a unique barcode at some initial time *t*_0_, and these barcodes are inherited by daughter cells with each cell division. After barcoding, the population develops over time and is then sampled at *t*_1_ and *t*_2_. Each sample is sequenced via scRNA-seq to determine the state (gene expression) of each cell as well as, if it is detected, the lineage barcode.

We are interested in how cell types with different growth rates are represented in multi-time clones, where the multi-time clones are constructed from sampling independently at *t*_1_ and *t*_2_ as illustrated by the example in Figure 1. Note that while we analyze a single pair of time points, the bias we discover may be greater for datasets with more than two time points (see Appendix A.2).

**Figure 1.**
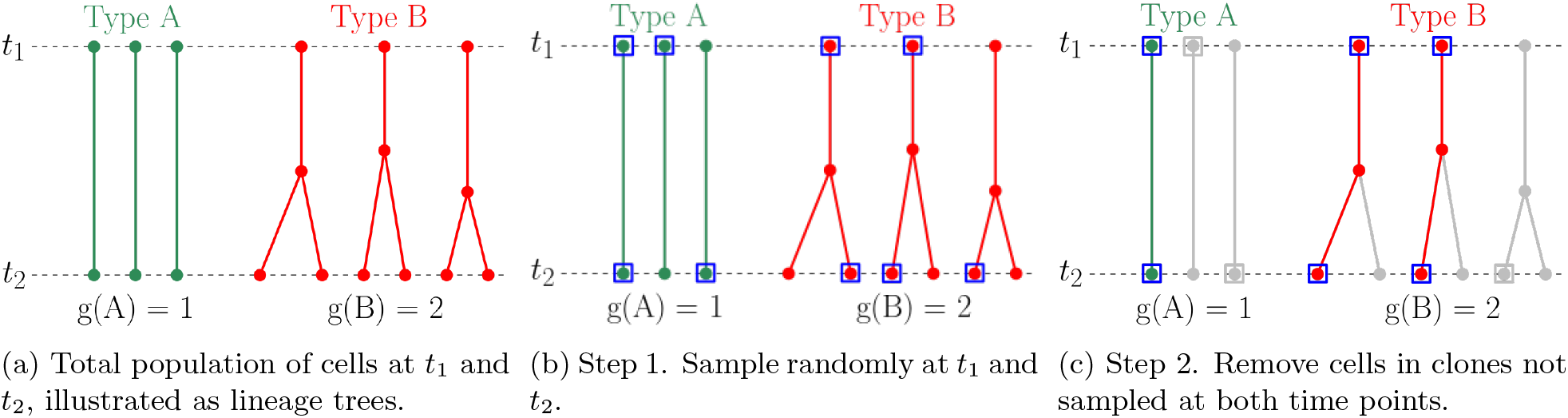
Example demonstrating the sampling process for obtaining the set of cells in multi-time clones: An illustration of the sampling process with two cell types, A and B, where A has growth 1, B has growth 2 and all lineages start with a single-cell at *t*_1_. The blue squares indicate cells sampled in step 1., and the grey lineages in (c) indicate the cells/lineages removed in step 2. The set of cells remaining in (c) are those in multi-time clones, denoted *MT*.

More precisely, for this derivation we will consider the set of cells in multi-time clones, denoted *MT*, which results from the sampling process as follows. Assume we have a population of 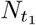 cells at *t*_1_ and 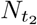 cells at *t*_2_. Each cell *x* at *t*_1_ gives rise to *g*(*x*) descendants at *t*_2_, as in the example shown in **Figure 1(a)**. Denote the set of cells in the population at *t*_1_ and *t*_2_ by ℳ(*t*_1_) and ℳ(*t*_2_) respectively. Using this notation, the sampling process may be described as:

1. Sample cells independently at *t*_1_ and *t*_2_, as illustrated in **Figure 1(b)**. Denote the sets of cells sampled by 𝒮(*t*_1_) ⊂ ℳ(*t*_1_) and 𝒮 (*t*_2_) ⊂ ℳ (*t*_2_).
2. Remove cells for which a lineage barcode was not detected from each sample. Denote the sets of remaining cells in each sample as ℬ(*t*_1_) ⊂ 𝒮(*t*_1_) and ℬ(*t*_2_) ⊂ 𝒮(*t*_2_). Remove cells in clones observed at only one time point (single-time clones), as in **Figure 1(c)**. We denote the remaining cells by *MT* ⊂ (ℬ(*t*_1_) ∪ ℬ(*t*_2_)). These are the cells in multi-time clones.

We now analyze the proportions of cell types in *MT* and find that these are different from the true abundances in the population at time *t*_1_. Similar derivations also show a bias in the abundances at *t*_2_ (see Appendix A.1), and, for time series with more than two time points, a bias in the abundances for each sample time (see Appendix A.2).

We analyze the proportions of cells at time *t*_1_ in MT by calculating the probability that a cell from the original population will be retained in MT. Denote the *sampling rate* (the proportion of cells sampled) by 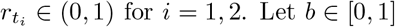 for *i* = 1, 2. Let *b* ∈ [0, 1] denote the probability that a lineage barcode is detected in a cell. We will refer to *b* as the *barcode rate*; it represents a combined rate of barcode insertion and detection. Denote the *net growth* between *t*_1_ and *t*_2_ of a single-cell *x* at *t*_1_ by *g*(*x*). This means that *g*(*x*) is the number of descendants of cell *x* at time *t*_2_, where *g*(*x*) = 0 if *x* dies between *t*_1_ and *t*_2_. We will show that **restricting to multi-time clones can bias the proportion of cells in a way that depends on the growth** *g*(*x*) **of the cells in the clone, sampling rates** 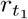 **and** 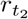, **and barcode rate** *b*.

We now derive an expression for the probability that a cell *x* at time *t*_1_ will be in *MT*. To simplify the expression of the probabilities, we assume that the two samples are independent and that we are sampling with replacement. Let 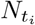 denote the number of cells in the population at time *t*_*i*_ for *i* = 1, 2. Define a clone membership function *C* by *C*(*x*) = *c* if cell *x* is in clone *c*. Finally, let 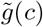 be the sum of the growth rates of the cells at *t*_1_ in clone *c*, i.e., 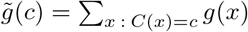. Then, for any cell *x* at time *t*_1_, the probability that *x* is retained in *MT* is

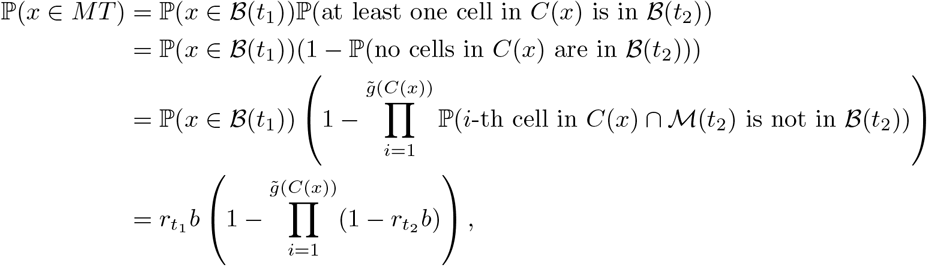

where the first line follows from our independent samples assumption, and the last line follows from approximating by sampling with replacement. Simplifying, we obtain

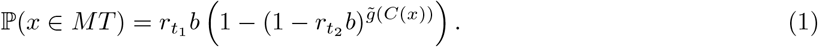

Equation 1 shows how the probability that a cell at *t*_1_ will be in a sampled multi-time clone depends on the growth rate of its clone relative to the other clones in the population at *t*_1_.

One may use Equation 1 to quantify the size of the bias effect on the proportions of cells from specific clusters in *MT* (e.g., cell types). See Equation 2 for an example of using Equation 1 to derive predicted cell type proportions in *MT*.

Next, we analyze how the proportions of cell types are impacted by this sampling bias. Consider the simplified setting in which: (1) each clone contains cells of only one cell type at *t*_1_, (2) the growth rate is homogeneous across each cell type (i.e. *g*(*x*) = *g*(*l*) for all cells *x* of type *l* at *t*_1_) and (3) all clones of type *l* start with the same number of cells at *t*_1_, denoted *m*(*l*). Under these simplifying assumptions, we have 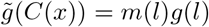 for every cell *x* of type *l*. Then the bias in proportions of cell types at *t*_1_ within *MT* is revealed by the following conditional probability,

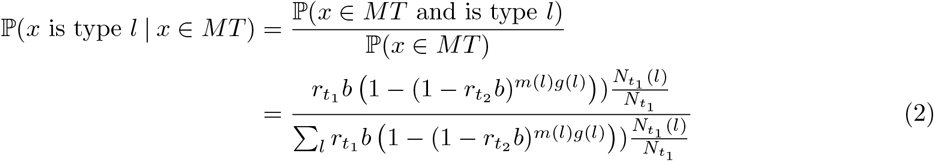

where the sum is over all possible cell types *l*. It is clear from Equation 2 that the proportions of the cell types at *t*_1_ in *MT* may significantly differ from the true proportions. For example, consider a population with two cell types *A* and *B* where *g*(*A*) = 1, *g*(*B*) = 2, *m*(*A*) = 1 and *m*(*B*) = 1. Suppose that the proportions of type *A* and *B* at *t*_1_ are equal, and at each time point we sample 50% of the population and detect a lineage barcode in every cell that we sample. Then, by Equation 2, the expected proportions of the two cell types *A* and *B* at *t*_1_ in *MT* are 2*/*5 and 3*/*5 respectively, in contrast to the 50:50 ratio of A:B that is present in the total sample at *t*_1_, as well as in the cells observed in single-time clones at *t*_1_. These proportions will depend on the growth rates, *g*(*l*) and *m*(*l*), of each cell type, the rates 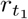 and 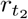 of cell capture, and the barcoding efficiency *b*.

The derivations in Appendix A.1 show that the cell type proportions at *t*_2_ in *MT* are similarly biased. However, at *t*_2_ the probabilities depend on growth rates between *t*_0_ and *t*_1_ rather than between *t*_0_ and *t*_2_ as in Equation 2. In this work, we focused our attention on the impact of the bias in the *t*_1_ proportions as a representative example of the effect.

To support our probabilistic reasoning, we conduct a simulation study to validate Equation 2. See Figure 2 for the results, where we plot the true simulated proportions and the proportions we predict using Equation 2. The plot shows the cell type proportions in *MT* for a population with two cell types, denoted *A* and *B*, with 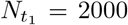, *m*(*A*) = 2 and *m*(*B*) = 4, *g*(*A*) = 1 and *g*(*B*) = 2, and varying sample rate *r* ∈ [2, 100%]. In this example, the true proportions of type *A* and *B* at *t*_1_ are equal. The plot clearly shows that the simulated results concentrate around the predictions and that the bias in the proportions increases at a superlinear rate as sampling rate *r* decreases, as expected from Equation 2. As sampling at each time is without replacement in these simulations, this provides evidence that approximating by sampling with replacement in the derivation of Equations 1 and 2 is reasonable for estimating the *t*_1_ cell type proportions in *MT*. The plot demonstrates that as sampling and barcode detection rates decrease, the bias in the cell type proportions in *MT* may diverge significantly from their true values.

**Figure 2.**
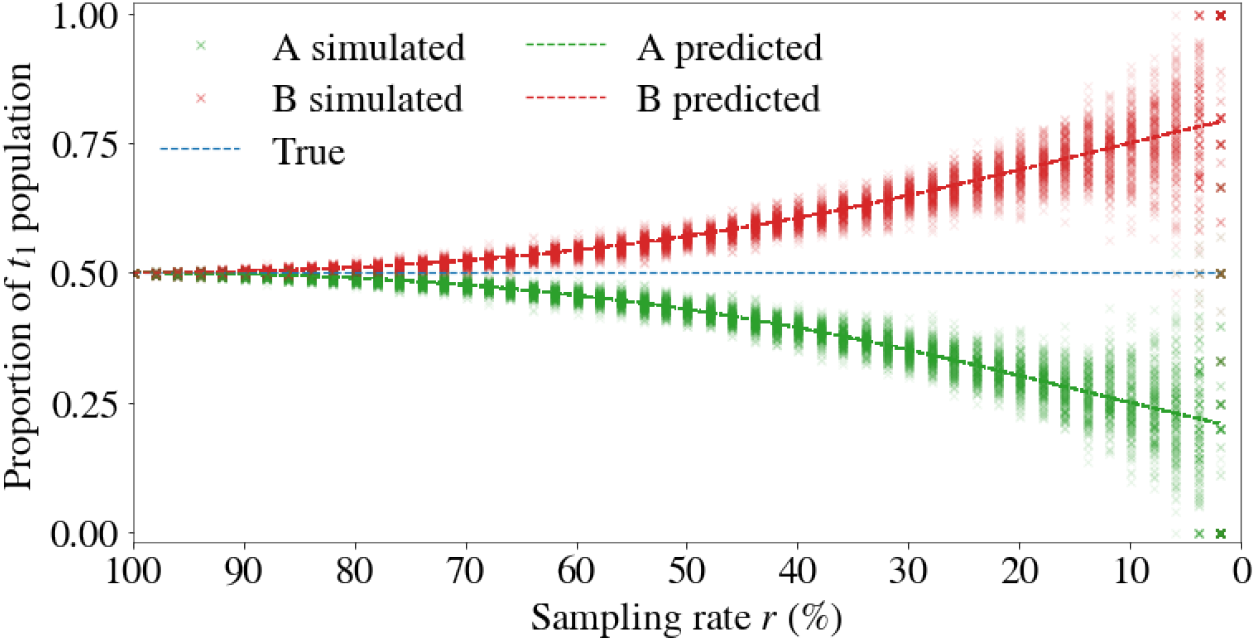
Restricting to multi-time clones introduces bias in cell type proportions. Simulated and predicted proportions of cell types A and B at *t*_1_ are shown as a function of the barcode sampling rate (combined sampling and barcode rate). The predicted proportions are computed from Equation 2 with 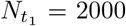 cells, *m*(*A*) = 2, *m*(*B*) = 4, *g*(*A*) = 1 and *g*(*B*) = 2. In this example, the true proportions of type *A* and *B* at *t*_1_ are equal. The bias in the proportions increases as sampling rate *r* decreases, where 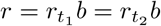 is plotted for 50 values between 100-2%. 200 replicates of the sampling process were generated for each of the 50 values.

Our investigations into this bias effect were motivated by a time course of statically barcoded blood cell development published in a separate paper [34]. We include here a summary (Figure 3) of the sampled cell type proportions in different subsets of clones: all (i.e., cells where a barcode was detected), multi-cell, and multi-time clones. *Multi-cell clones* are those that contain more than a single cell.

**Figure 3.**
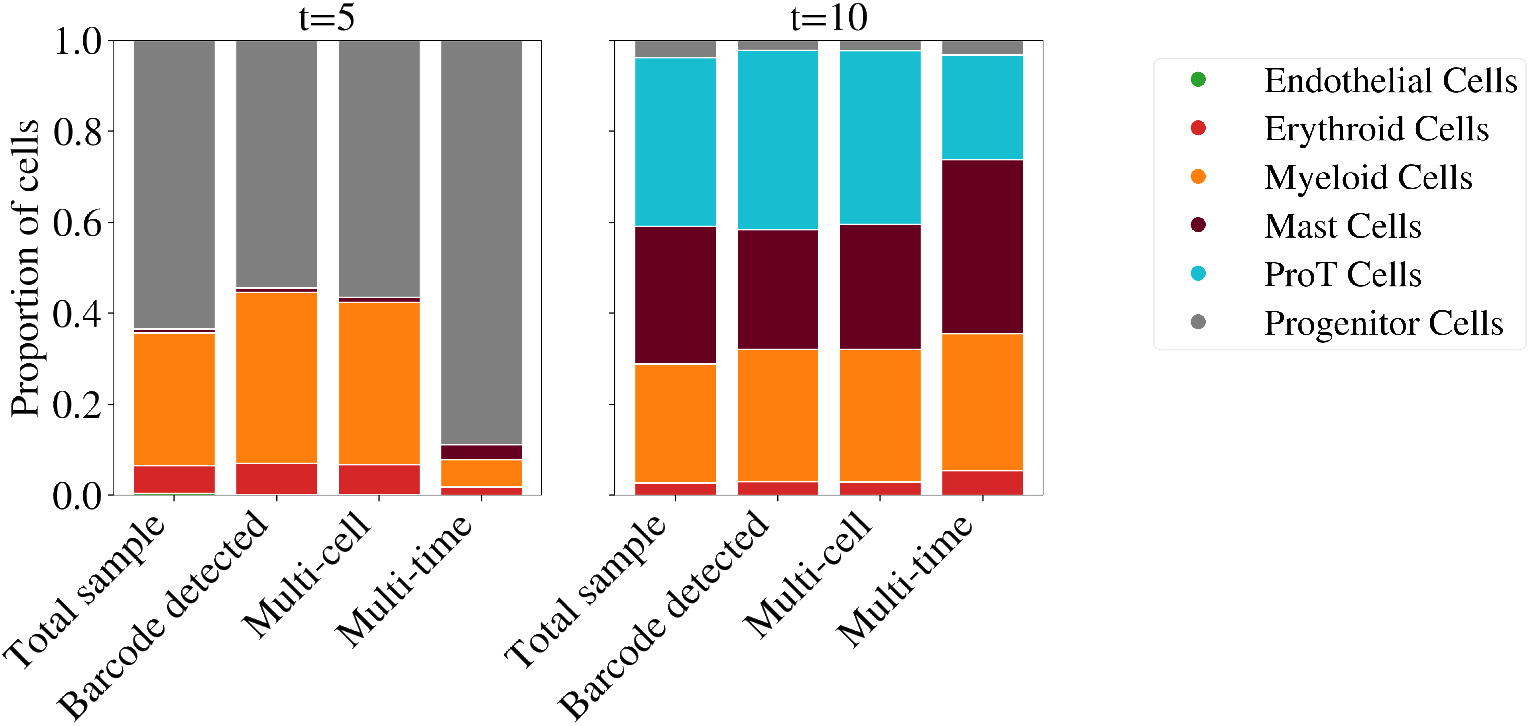
Sampled cell type proportions in a time course of statically barcoded blood cell development differ when restricting to the subset of cells in multi-time clones. The proportions of different cell types observed at two time points (*t* = 5 and *t* = 10) in different subsets of clones (barcode detected, multi-cell and multi-time) are shown as coloured areas in bar plots. The proportions at each time for the total sample, barcode detected and multi-cell clones are similar (variance within that expected from random sampling), whereas the cell type proportions in the multi-time clones are significantly different at both time points. In particular, at *t* = 5, the proportion of progenitor cells is approximately 0.9 instead of being in the range 0.5-0.7, and the myeloid proportion is approximately 0.1 instead of being in the range 0.3-0.4. The difference for the proportions at *t* = 10 is smaller, but the proportions in the multi-time clones are still the furthest from those of the total sample.

From our derivations, we do not expect biased cell type proportions in the set of cells where a barcode was detected or those in a multi-cell clone. Indeed, in Figure 3 the difference in proportions between the total sample and these two subsets is within what one would expect from unbiased random sampling. In contrast, the difference between the proportions in the total sample and those in multi-time clones is outside the expected variance of unbiased sampling. The bias we have identified may explain this difference and be driven by a higher relative growth rate of progenitor cells. Conclusively determining that this is the cause would require growth and death rate estimates that were not available for this dataset. This rate information is needed to eliminate the alternative explanation that the lineages of cell types with decreased proportions (e.g., myeloid at *t* = 5) died out by *t* = 10 at a significantly greater frequency than those of increased proportions (e.g., progenitor at *t* = 5).

We conclude with four summary remarks related to this statistical phenomenon: (1) single-time lineage information exhibits no such bias, (2) there will be no bias when growth rates are equal across the population for all time, (3) the bias can present for clones sampled at just two time points, but it may increase with sampling at additional times (see Appendix A.2) and, (4) the cell type proportions in *MT* at each sample time may be biased depending on the growth rates in a similar manner as in Equation 2. These remarks and the results of this section apply to the proportions of any subdivision/clustering of cells into subpopulations that correlate with growth rates, cell types being one prominent example.

## 3 Bias in multi-time clonal barcodes can affect trajectory inference

Given this newly discovered sampling bias in multi-time clonal barcodes, a natural question is whether this bias impacts the current state-of-the-art methods which use this information for trajectory inference.

We investigate the answer to this question for *CoSpar*, a method that has shown promising results for datasets representing hematopoiesis, reprogramming and directed differentiation [31]. *CoSpar* (coherent, sparse optimization) features two different methods for lineage-informed trajectory inference. Both methods use cell state information, but one uses only single-time clonal data while the other uses only multi-time clonal data. We will refer to the latter as *CoSpar-MT*. The method is built on two key assumptions about trajectory couplings: (1) the coupling is ‘locally coherent’, and (2) the coupling is sparse. These two assumptions are built into the *CoSpar* method via a ‘smoothing’ and a thresholding step respectively [31].

We focus on the impact of the bias on the inferred fate probabilities of the cells, a key prediction derived from trajectory couplings. Our results demonstrate that it is possible for the bias in the multi-time clone data to negatively impact the performance of *CoSpar-MT* on the task of fate probability prediction (see Figure 4). The results were gathered for a simple simulated population with two cell types at *t*_1_ and *t*_2_ for which the ground truth fate probabilities are known.

**Figure 4.**
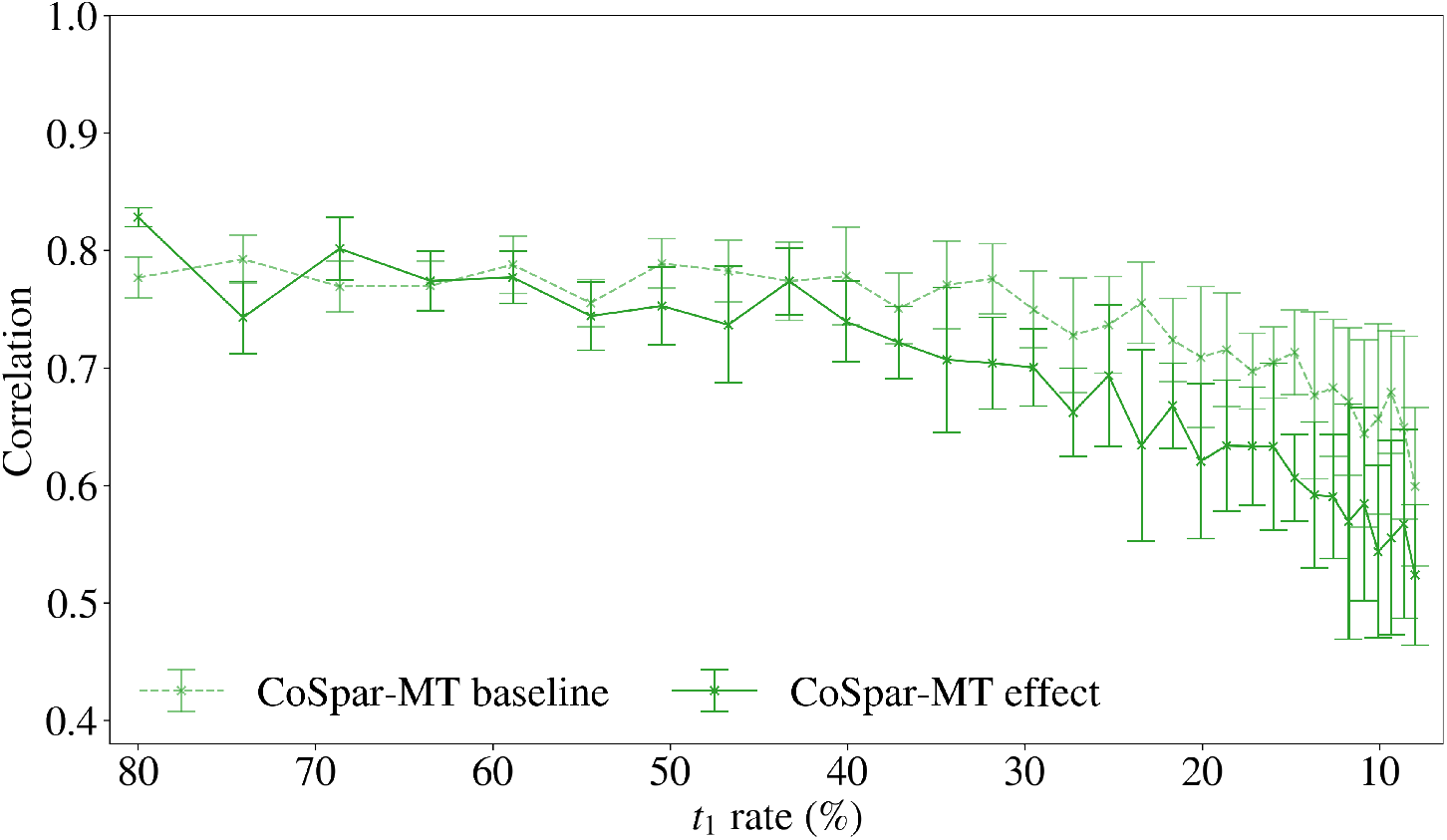
It is possible for the bias to impact fate probability predictions from CoSpar-MT: Pearson correlation coefficient between the true fate probability and the fate probability estimated from *CoSpar-MT* (the probability that each cell is fated to be type A at *t*_2_) as a function of *t*_1_ rate (combined sampling and barcode rate 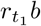) for *b* = 0.8 fixed for the *baseline* and *effect* cases. Values are summarized as a mean across ten replicates with error bars given by the standard deviation. The plot shows that the mean correlation decreases as the sampling rate decreases and that once the bias effect is sufficiently large (around 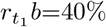) the mean correlation is lower in the *effect* case than the *baseline* case with a difference of approximately 0.1. This demonstrates that it is possible for the bias in multi-time clones to negatively impact fate probability prediction. The results for fate B are identical and hence are omitted.

Two variations, *baseline* and *effect*, of the population were simulated to separate the effect of the bias in the multi-time clones from the effect of the sampling rate. In the *baseline*, all growth rates are equal resulting in no bias effect. In the *effect* case, the growth rates vary between two and eight descendants across the population, resulting in a bias in the *t*_1_ cell type proportions in the multi-time clones. The growth rate distribution includes a mixture of higher and lower growth cells in one of the cell types, which could represent cells with a distinct chromatin accessibility profile (between *t*_1_ and *t*_2_) to the rest of the cluster.

To control for the effect of the number of cells sampled and manage the computational resources required for trajectory inference, 1000 cells were sampled at *t*_1_ and *t*_2_ for every sampling rate. Sampling a fixed number of cells rather than sampling at a fixed rate also more realistically models experimental procedures, in which only a fixed budget of cells can be sequenced. Each specified *t*_1_ sampling rate, 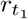, is obtained by simulating a population of 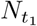 cells where 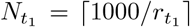. For the *effect* and *baseline* cases each, the results were generated from 30 simulated populations (one for each *t*_1_ sampling rate tested), each containing between 2500 and 60 000 cells with population size scaling inversely to sampling rate. The true trajectory coupling was computed for each population, and then 1000 cells were sampled at each *t*_1_ and *t*_2_. Next, *CoSpar-MT* was used to estimate the trajectory coupling between the *t*_1_ and *t*_2_ samples.

The results in Figure 4 show that the mean correlation decreases as the sampling rate decreases, and furthermore, once the bias effect is sufficiently large (around 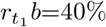) the mean correlation is lower in the *effect* case than the *baseline* case, with a difference of approximately 0.1. Since the error bars of the *baseline* and *effect* results overlap, we do not claim that correlation is lower in the *effect* case with high probability, but the trend of the mean is clear. This demonstrates that it is possible for the bias in multi-time clones to negatively impact fate probability predictions from state-of-the-art methods.

## 4 *LineageOT-MT* : Extension of LineageOT to use multi-time clonal data

*LineageOT* [1] is an extension of the state-based *WaddingtonOT* framework [12] to incorporate lineage information from single-time lineage trees (trees where all observed cells are from a single time point). The current *LineageOT* method neglects to use valuable lineage information encoded in multi-time clonal barcodes, as is available in static lineage barcoding with population subsampling across time points. This limitation motivated our extension of *LineageOT* to use *both* multi-time and single-time clonal barcode information. We refer to this extension as *LineageOT-MT*^1^. Our approach in *LineageOT-MT* differs from the approach in *CoSpar* where either multi-time (as in *CoSpar-MT*) *or* single-time clonal barcodes are used but *not both*.

The extension requires no new mathematical theory and only a relatively small change in the implementation of the algorithms. We simply condition on the state of all observed cells in each clone in the ancestor estimation step in *LineageOT*, rather than just those at the later time point. This estimation is performed by modeling the lineage trees as Gaussian graphical models relating the cell states [1]. In the extension, we condition in the Gaussian model on all cells observed in the lineage tree (over all time points), not just those sampled from the later time point being coupled. The Gaussian estimation now benefits from using the observed states of ancestral and present-time clonal relatives in addition to the descendant relatives.

Trajectory inference results from simulated datasets show that multi-time clonal information improves the performance of *LineageOT*. Figure 5 demonstrates the improvement possible for the task of fate probability prediction. The results from this figure describe prediction performance on a simulation scenario similar to those used in the analysis of *CoSpar* in Section 3. Additional such comparative results were gathered for growth rate variations of this scenario and the simulation scenarios described in Section 3, using over 1200 simulated datasets in total (ten replicates per each of the 120 variations). *LineageOT-MT* achieved an equal or higher mean correlation with the true fate probability than *LineageOT* on all of these datasets.

**Figure 5.**
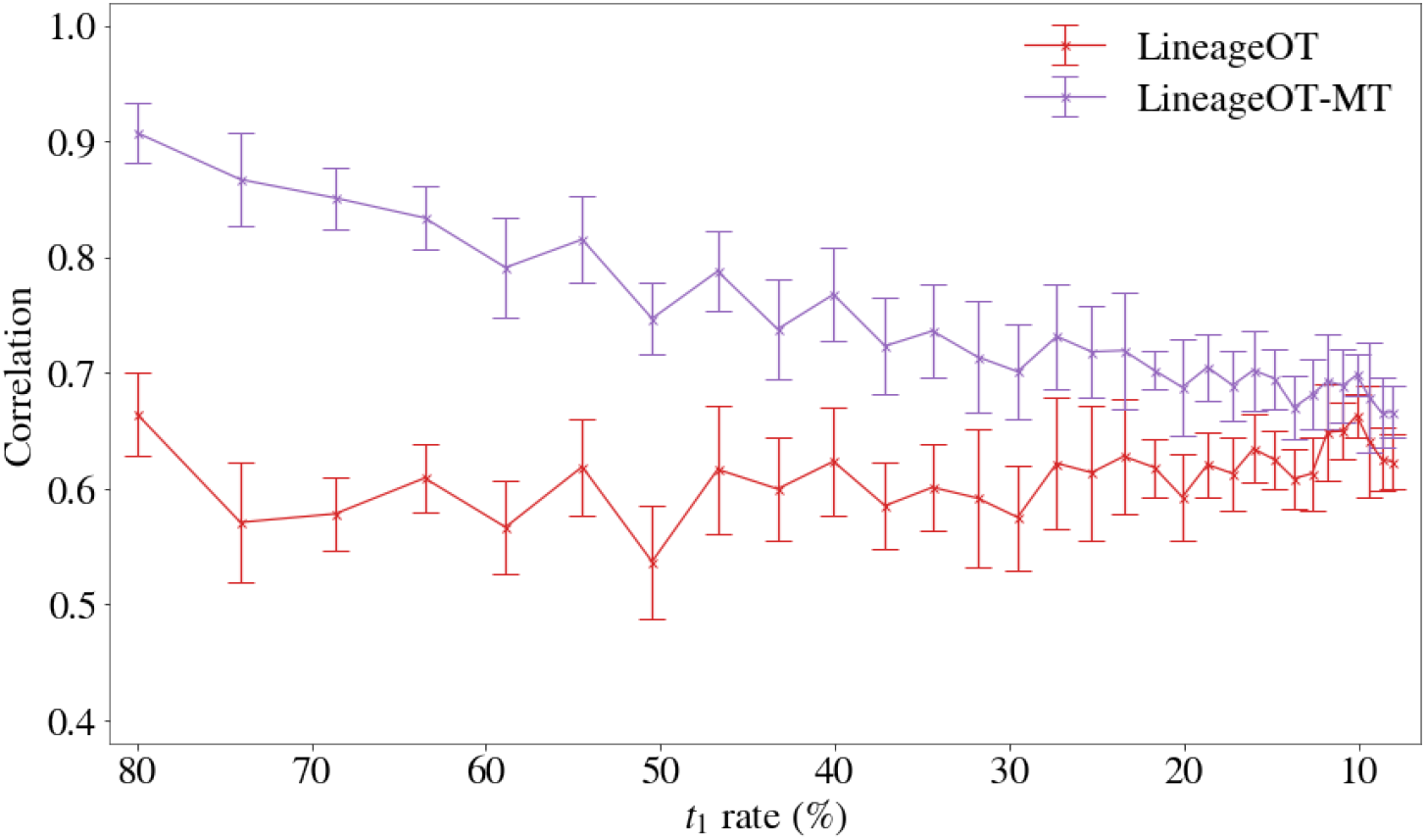
Multi-time information improves the fate probability predictions from LineageOT: Pearson correlation between the true fate probability and the fate probability estimated from *LineageOT* and *LineageOT-MT* for fate A at *t*_2_ as a function of *t*_1_ rate (combined sampling and barcoding rate 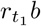) for *b* = 0.8. Values are summarized as a mean across ten replicates with error bars given by the standard deviation. The results for fate B are identical and hence are omitted.

The results in Figure 5 show that at the highest rate tested of 80% our extension of the method increases correlation with the true fate probability from approximately 0.6 to over 0.9. The performance of *LineageOT-MT* over *LineageOT* was assessed on simulated datasets across a range of sampling rates, as shown in Figure 5, to illustrate the effect of decreasing the available multi-time clonal barcode information on prediction performance. As the sampling rate decreases, the decline in performance of *LineageOT-MT* is approximately linear and converges towards the performance of *LineageOT*. This is expected since the amount of multi-time clonal information is decreasing to zero with the sampling rate.

*LineageOT-MT* incorporates additional information about the developmental process compared to *LineageOT*, so these results are not surprising. However, the size of the boost in performance is encouraging for further development in this direction.

### 4.1 Effect of bias in multi-time clonal barcodes on *LineageOT-MT*

In this section, we explore the impact of multi-time clonal barcode bias on *LineageOT-MT*. This exploration is analogous to the study of the effect of the bias on *CoSpar-MT* in Section 3.

The results were gathered using *baseline* and *effect* variations (as defined in Section 3) of the simulation scenario that was used to benchmark the performance of *LineageOT-MT* above. The performance of *LineageOT-MT* over decreasing sample rates for these two variations is shown in Figure 6. This figure demonstrates that the mean correlation decreases as the sampling rate decreases. Furthermore, for a relatively high range of sampling rates (from *t*_1_*b* =0.8 to around *t*_1_*b* =0.3), the mean correlation is lower in the *effect* case than in the *baseline* case, with a difference of approximately 0.1. These results demonstrate that it is possible for the bias in multi-time clones to negatively impact fate probability predictions from *LineageOT-MT*.

**Figure 6.**
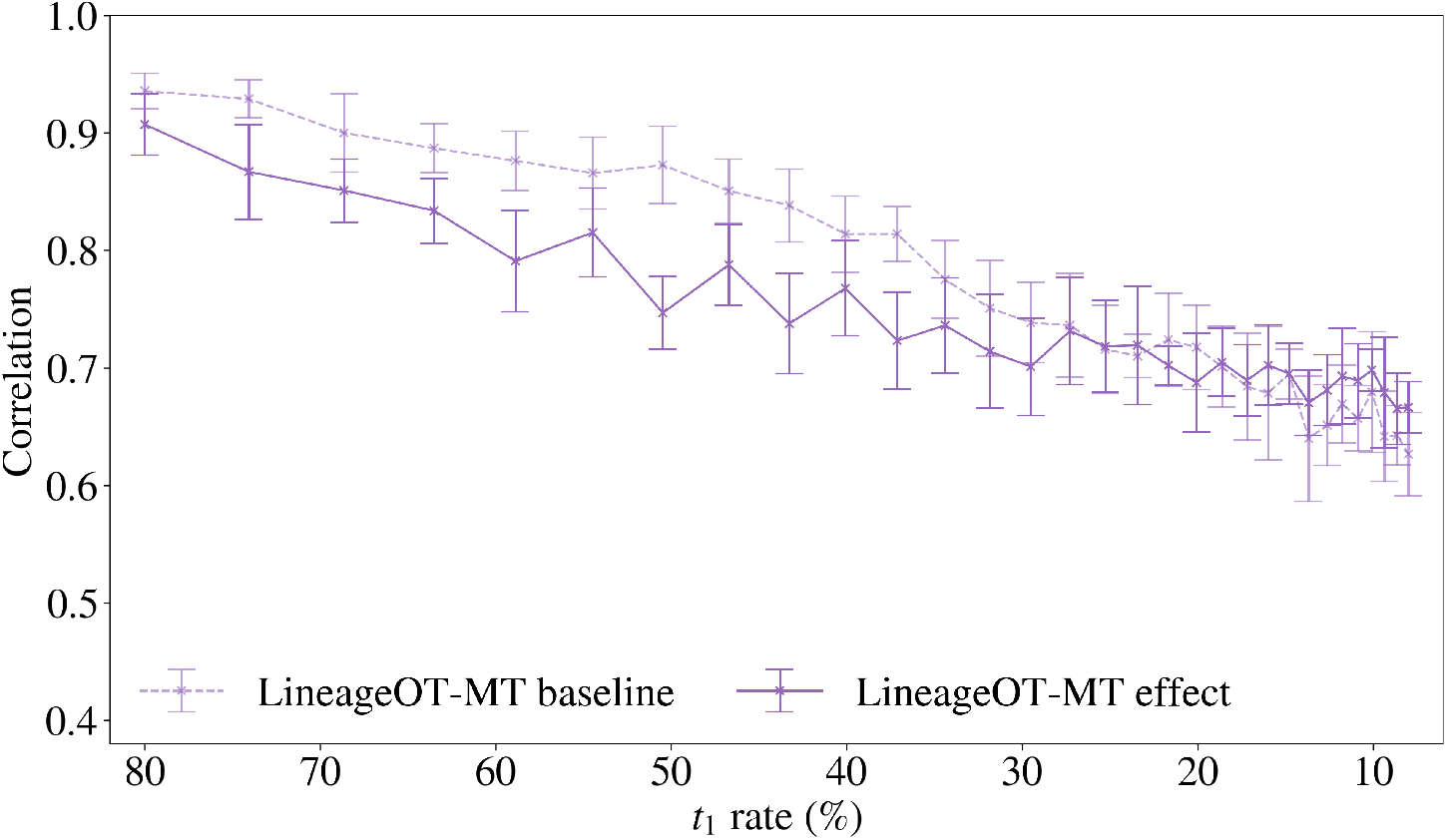
It is possible for the bias to impact fate probability predictions from LineageOT-MT: Pearson correlation between the true fate probability and the fate probability estimated from *LineageOT-MT* (the probability each cell is fated to be type A at *t*_2_) as a function of *t*_1_ rate (combined sampling and barcode rate 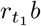) for *b* = 0.8 fixed for the *baseline* and *effect* cases. Values are summarized as a mean across 10 replicates with error bars given by the standard deviation. The plot shows that the mean correlation decreases as the sampling rate decreases, and that for sufficiently *high* sampling rate (greater than approximately 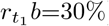) the mean correlation is lower in the *effect* case than the *baseline* case, with a difference of approximately 0.1. This demonstrates that it is possible for the bias in multi-time clones to negatively impact fate probability prediction. The results for fate B are identical and hence are omitted.

The lack of impact on *LineageOT-MT* at rates less than 0.3, where the size of the bias effect is larger, may be explained by the methodological ‘convergence’ of *LineageOT-MT* to *LineageOT* as the sampling rate decreases. This convergence corresponds to a shift from using *multi-time* and *single-time* lineage information to only using *single-time* lineage information. The number of cells observed in multi-time clones decreases with the sampling rate at a faster rate than the number in single-time clones, resulting in this shift occurring before the sampling rate nears zero. Recalling that the single-time information is not biased, this convergence may counter the impact of the bias on cell fate predictions from *LineageOT-MT*.

## 5 Discussion

In this work, we consider the problem of developmental trajectory inference using lineage information derived from multi-time clonal barcodes. We have shown that in certain experimental settings, there may exist a bias in the proportions of cell types represented in the multi-time clonal barcode data. This sampling bias emerges for clones sampled at just two time points and may increase with sampling at additional times. The experimental settings where this bias emerges are those with heterogeneous growth rates and a combined sampling and barcode detection rate of less than 100% for at least one sample in the time course. These are of course very general conditions, and we suspect this bias to be present in most multi-time clonal barcode datasets.

In addition, we showed that it is possible for this bias to negatively impact predictions made from inferred developmental cell trajectories. This was demonstrated on a state-of-the-art trajectory inference method, *CoSpar* [31], using simulated datasets for which the ground truth is known.

Our third contribution is an extension of the *LineageOT* trajectory inference method [1] to additionally use multi-time clonal barcode information. This extension, *LineageOT-MT*, robustly improves the performance of the method. We then showed that it was possible for the bias to impact predictions made from trajectories inferred using *LineageOT-MT*. We note that there may be more sophisticated ways to integrate the multi-time clone data into the *LineageOT* method beyond our present approach that could yield further performance improvements and robustness.

Our studies on the impact of the sampling bias on fate probability prediction with *CoSpar-MT* and *LineageOT-MT* show that each method is sensitive to the bias for distinct scenarios. Certain features of each method mediate the robustness or sensitivity to the bias effect. Qualitatively, the smoothing step in *CoSpar-MT* (which is strongly dependent on the state space geometry of the sampled cells) seems to improve the robustness of the method to the bias effect in some settings. For *LineageOT-MT*, using cell state and single-time clonal barcode information in addition to multi-time appears to provide robustness to the bias effect. This conclusion is supported by these first two data sources being unbiased and the prediction accuracy of the method being unaffected in the low sampling rate regime. Purely considering the trends observed with respect to sampling rate, *LineageOT-MT* may offer robustness to the bias effect in the lower sampling rate regime while *CoSpar* may offer robustness at higher rates.

Multi-time clonal data, even if known to be biased by this sampling effect, is too valuable not to leverage in the study of single-cell development. This point is evidenced in this paper by the robust improvement of *LineageOT-MT* over the original *LineageOT*. What is needed is a solution for correcting the bias. Our work suggests that the most promising avenue is to leverage the unbiased single-time clonal barcode and cell state data.

We observe that the smoothing step of the *CoSpar* algorithm could be considered a kind of label propagation or ‘pseudo-labeling’ [35]–[37] that leverages the unbiased cell state data. The smoothing effectively propagates the multi-time clone label information from cells to nearby cells (in state space) that were not observed in a multi-time clone. See Figure 7 for an illustration of the effect of label propagation on a dataset. Label propagation is one form of semi-supervised learning that can be applied to barcoded single-cell datasets. *Semi-supervised learning* refers to inference in settings where only a subset of the training data is labeled. This is the case in our application, where some cells are labeled (by their clone) and some are unlabeled as a lineage barcode was not detected. For any label propagation approach, it will be important to account for growth rates in the propagation to generate labels that correct the bias in the multi-time clones.

**Figure 7.**
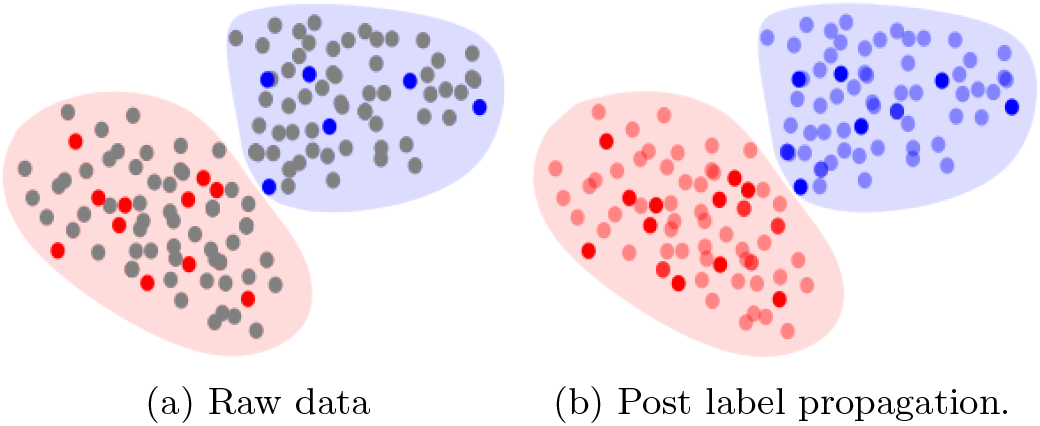
An illustration of label propagation: The left image illustrates a raw partially labeled dataset where the dots are data points coloured by their label. The grey dots are the unlabeled points and the background colour shows the true distribution of each cluster. The right image shows the same dataset after performing successful label propagation to share labels from the labeled points to the unlabeled points, resulting in a fully labeled dataset.

Motivated by the concept of pseudo-labeling and the unbiased nature of single-time clonal data, one idea to improve the robustness of *CoSpar-MT* to the bias is to correct the multi-time clonal matrix in the algorithm [31] using iterative proportional fitting (IPF). IPF is also known as Sinkhorn’s algorithm in the optimal transport literature. It is a procedure for fitting data to a set of marginals which are typically known to be more accurate than the those of the data itself. In the case of *CoSpar-MT*, the two marginals of the multi-time clonal matrix correspond to the proportion of clones present in each subpopulation at *t*1 and *t*_2_. The idea is to use IPF to project the matrix to the subspace of matrices with the “correct” marginals, i.e. those given by the unbiased single-time clonal data. Further investigation is required to determine whether this correction would indeed eliminate the impact of the bias in any predictions made using *CoSpar-MT*.

The equations in Section 2, and studies on *CoSpar-MT* and *LineageOT-MT* in the subsequent sections, provide some indication of when the bias effect will impact trajectory inference. In the future, it would be useful to have a statistical test that quantifies the expected impact of the bias on trajectory inference. A practical statistical test should use only information available in standard datasets of lineage barcoded sequencing time courses, which importantly may not include precise estimates of cell growth rates. An-other important step is the definitive identification of the bias effect and its impact on trajectory inference methods for a real dataset. We leave such developments to future work, having focused in this work on the mathematical details and validations only possible using simulated data.

We hope that our identification of this statistical phenomenon serves to inform practitioners using multi-time clonal barcodes of possible biases in the data, and guide future method development. Beyond their use in proving the existence of the bias, the probabilities derived in Section 2 can also be used as a tool by practitioners for approximating the size of the bias effect in a dataset. Our contributions beyond these points are our extension to *LineageOT*, and the above discussion of how methods might be developed to overcome the impact of this sampling bias. Such development is an important endeavour to allow practitioners to make use of the valuable information in multi-time clonal barcodes, without these concerns of biased results and conclusions.

## A Appendix

### A.1 Derivation of biased proportions in *MT* at *t*2

We include below the derivation of probabilities of subpopulation proportions at *t*_2_, which is analogous to the derivation included in the paper for *t*_1_ (Equations 1 and 2). Derivations for time points beyond *t*_1_ and *t*_2_ would follow similar arguments.

As we derive below, the cell type proportions in *MT* (as defined in Section 2) at *t*_2_ are biased by the relative cell growth between *t*_0_ and *t*_1_. Let *m*(*c*) denote the number of cells in clone *c* at *t*_1_, or equivalently, the net growth rate between *t*_0_ and *t*_1_ of the cell at *t*_0_ which was barcoded as clone *c*. Then we have:

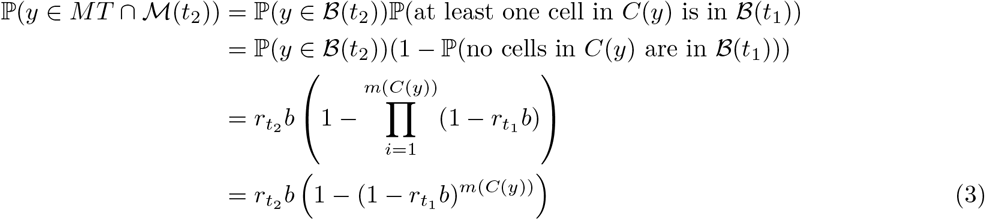

and, if we simplify the expression by assuming that all clones of type *l* have the same value of *m*(·), which we denote *m*(*l*) as in the derivation of Equation 2, then we have

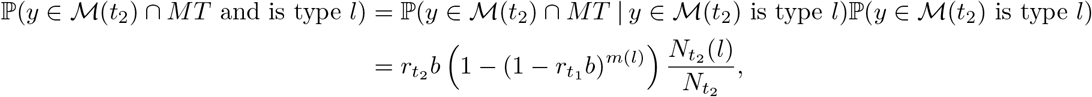

and therefore we obtain,

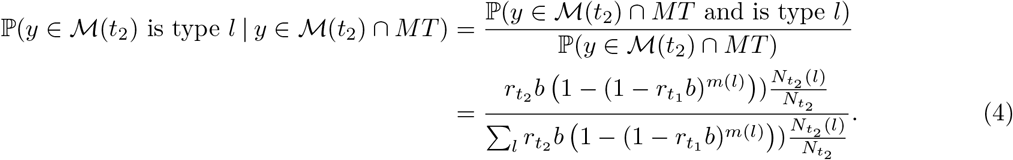

These equations are very similar to those for *t*_1_ (Equations 1 and 2) except here the probabilities depend on growth rates between *t*_0_ and *t*_1_ rather than between *t*_0_ and *t*_2_. Comparing Equation 4 to Equation 2, if the true cell type proportions at *t*_1_ and *t*_2_ are the equal then the bias in the proportions may be lesser or greater at *t*_1_ than at *t*_2_ (depending if *g*(*l*) *>* 1.0 or *<* 1.0 respectively) due to the exponent being *m*(*l*)*g*(*l*) rather than *m*(*l*).

### A.2 Derivation of biased proportions in *MT* for an arbitrary number of time points

We include below the derivation of the probabilities of subpopulation proportions at any sample time *t*_*k*_ ∈ {*t*_1_, *t*_2_, …, *t*_*T*_} in a time course with a finite but arbitrary number of time points *T* ≥ 2. This derivation follows a similar argument to that for *t*_1_ and *t*_2_ for *T* = 2 in Section 2 and Appendix A.1 respectively.

Let *x* denote a cell alive at time *t*_*k*_ and let *r*_*t*_ denote the sampling rate at time *t* ∈ {*t*_1_, *t*_2_, …, *t*_*T*_}. Then, by the law of total probability and De Morgan’s law, we have:

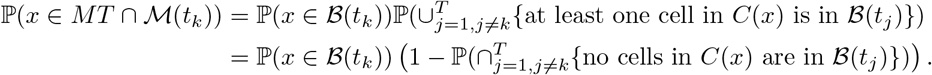

Now let 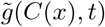 denote the sum of the growth rates of all cells alive in clone *C*(*x*) at time *t*. Then, by the same independence assumption made in Section 2, the probability of the intersection can be broken down as the following product,

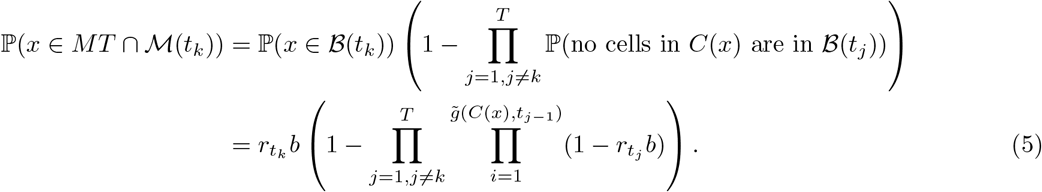

By Equation 5, since each factor in bounded in [0, 1] (as they represent probabilities), as *T* increases the product over the time points will be non-increasing, and hence the size of ℙ(*x* ∈ *MT ∩ ℳ*(*t*_*k*_)) will be non-decreasing. To reason how this effects the biased proportions of subpopulations we consider the following probability for some cell type (or any subpopulation correlated with growth) *l*:

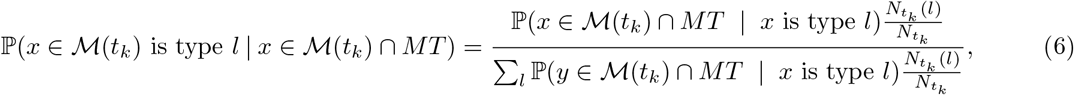

where ℙ(*y* ∈ ℳ(*t*_*k*_) ∩ *MT* | *x* is type *l*) takes an analogous form to in Section 2 based off of Equation 5. From Equation 6 and the observation that ℙ(*x* ∈ *MT* ∩ ℳ(*t*_*k*_)) will be non-decreasing as *T* increases, the behaviour of the probability representing the biased proportion ℙ(*x* ∈ ℳ(*t*_*k*_) is type *l* | *x* ∈ ℳ(*t*_*k*_) ∩ *MT*) as *T* increases will depend on how the denominator changes. More specifically, since 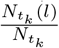 is constant for all *l* as *T* increases, it will depend on how the values of P(*x* ∈ M(*t*_*k*_) ∩ *MT* | *x* is type *l*) behave for all types *l*.

Let us consider the case in which ℙ(*x* ∈ *ℳ*(*t*_*k*_) ∩ *MT* | *x* is type *l*) only increases for type *l* (for all other types the probability stays constant as *T* increases). Then it is easy to prove with simple algebraic manipulation that the new expression (after the increase in *T*) on the right hand side of Equation 6 is greater than the previous expression. This case demonstrates that the size of the bias effect may increase as *T* increases beyond *T* = 2. In general, whether or not the bias effect will increase depends on complex interactions between the growth rates in the new periods of time in the time course, as summarized with Equation 6. This equation may be used to estimate the size of the bias effect as demonstrated using Equation 2 in Section 2.

## Footnotes

1 The implementation of *LineageOT-MT* is publicly available on a fork of the *LineageOT* package at: https://github.com/rbonhamcarter/LineageOT/tree/multi-time-clones.

